# Dependence of aerosol-borne influenza A virus infectivity on relative humidity and aerosol composition

**DOI:** 10.1101/2024.05.28.596202

**Authors:** Ghislain Motos, Aline Schaub, Shannon C. David, Laura Costa, Céline Terrettaz, Christos Kaltsonoudis, Irina Glas, Liviana K. Klein, Nir Bluvshtein, Beiping Luo, Kalliopi Violaki, Marie O. Pohl, Walter Hugentobler, Ulrich K. Krieger, Spyros N. Pandis, Silke Stertz, Thomas Peter, Tamar Kohn, Athanasios Nenes

## Abstract

We describe a novel biosafety aerosol chamber equipped with state-of-the-art instrumentation for bubble-bursting aerosol generation, size distribution measurement, and condensation-growth collection to minimize sampling artifacts when measuring virus infectivity in aerosol particles. Using this facility, we investigated the effect of relative humidity (RH) in very clean air without trace gases (except ∼400 ppm CO_2_) on the preservation of influenza A virus (IAV) infectivity in saline aerosol particles. We characterized infectivity in terms of 99%-inactivation time, *t*_99_, a metric we consider most relevant to airborne virus transmission. The viruses remained infectious for a long time, namely *t*_99_ > 5 h, if RH < 30% and the particles effloresced. Under intermediate conditions of humidity (40% < RH < 70%), the loss of infectivity was the most rapid (*t*_99_ ≈ 15-20 min, and up to *t*_99_ ≈ 35 min at 95% RH). This is more than an order of magnitude faster than suggested by many previous studies of aerosol-borne IAV, possibly due to the use of matrices containing organic molecules, such as proteins, with protective effects for the virus. We tested this hypothesis by adding sucrose to our aerosolization medium and, indeed, observed protection of IAV at intermediate RH (55 %). Interestingly, the *t*_99_ of our measurements are also systematically lower than those in 1-μL droplet measurements of organic-free saline solutions, which cannot be explained by particle size effects alone.

## 1. Introduction

Respiratory diseases are transmitted by exhaled aerosol particles over a wide spectrum of particle sizes produced by speaking, coughing, sneezing as well as forced and rest breathing. Previously considered unimportant for virus transmission (e.g., Randall et al., 2021), virus-containing aerosol particles are now thought to be a major vehicle for spreading respiratory diseases, including influenza and COVID-19 (Coronavirus disease 2019; Chen et al., 2020); Greenhalgh et al., 2021; Miller et al., 2021). Numerous epidemiological studies have pointed to the importance of environmental conditions on the stability of airborne viruses (e.g., Hanley and Borup, 2010; Deyle et al., 2016; Sehra et al., 2020). It is critical that laboratory experiments inform transmission modelling and epidemiological studies to gain new insights into how environmental factors influence virus infectivity, and how they can be used to mitigate virus transmission.

The most common approaches for studying airborne virus infectivity include the injection of virus-containing particles into a flow-thru chamber, a rotating drum or a static chamber, as well as single-particle levitation to expose the viruses to controlled conditions and then sample them for characterization of their infectivity or other properties. The approaches adopted differ in chamber volume, particle formation, suspension time, suspension medium and experiment duration according to the needs of the experiment (Santarpia et al., 2020; Le Sage et al., 2023). Although these methods are established, they still lack standardization, rendering the reproducibility and comparability of infectivity studies difficult (Santarpia et al., 2020; Yeh and Setser, 2022).

Several factors are known to contribute to the inactivation of an airborne virus, including physico-chemical parameters of the particle, relative humidity (RH), air temperature and composition of the surrounding gas. The influence of air temperature has been extensively documented based on exposure experiments and is now relatively well characterized (e.g., Harper, 1961; Lowen et al., 2007 and Gustin et al., 2015). Experiments in which virus-containing aerosol particles were exposed to controlled levels of relative or absolute humidity were initiated as early as the 1940s (e.g., Loosli et al., 1943 and Edward et al., 1943 with animal models) although there is still no consensus concerning the effect of this environmental factor on virus infectivity. Conversely, the impact of vapor partitioning of CO_2_ and of acidic gases in the ambient air on particle pH and subsequent virus inactivation has only recently become a topic of scientific interest and is still widely debated (Yang and Marr, 2012; Oswin et al., 2022a; Luo et al., 2022; Klein et al., 2022; Oswin et al., 2022b).

Experiments that investigated the dependence of airborne virus infectivity on relative or absolute humidity have been reviewed by Sobsey and Meschke (2003), Yang and Marr (2012), and Božič and Kanduč (2021). Such experiments showed that enveloped viruses (e.g., coronaviruses, bacteriophage φ6, coliphage and various influenza virus strains) tend to better retain their infectivity at low RH, in contrast to non-enveloped viruses for which a high RH is favorable. Several exceptions to these findings, however, remain unexplained and conflicting results are found even for a single type of virus. For influenza A virus (IAV), some studies reported an infectivity versus RH dependence that follows a “U-shape” or “V-shape”, with inactivation most rapid at intermediate RH between 40% and 70% (Shechmeister, 1950; Schaffer et al., 1976; and Gustin et al., 2015 using animal models). Others found a slowing of inactivation from low to intermediate RH, but little response with further RH increase (Harper, 1961; Hemmes et al., 1962). Yang et al. (2012) tried to explain these disagreements and hypothesized, based on experiments with 1-μL deposited droplets, that the protein-to-salt concentration ratio might protect IAV at intermediate RH. Recent studies (Kormuth et al., 2018; Dubuis et al., 2021) further support these findings. Yet, the mechanism behind such a protective effect remains poorly understood.

The interpretation of these partially contradictory results is complicated by differences in the techniques used in each study. A majority of airborne virus exposure experiments (e.g., Benbough, 1971; Donaldson and Ferris, 1976; Karim et al., 1985; van Doremalen et al., 2013; Verreault et al., 2015; Pyankov et al., 2018; Dubuis et al., 2021) used a Collison nebulizer (CH Technologies, Westwood, NJ, USA) and a glass impinger such as the BioSampler (SKC, Eighty Four, PA, USA). The Collison produces small particles by pulsing high-velocity air through a small orifice while aspirating the liquid suspension, and projecting this stream onto the glass sides of the jar. Larger particles are trapped and recirculated. While this twin-fluid fragmentation technique promotes high reproducibility and particle output, concerns have emerged regarding the effects of impaction, shear forces and recirculation on virus infectivity, possibly damaging the virus structure in a way that does not occur during exhalation in infected people. Similar forces and consequences might be encountered when sampling with the BioSampler, which uses nozzles to tangentially impact airborne particles onto a thin layer of liquid formed on a glass surface. On the contrary, the SLAG nebulizer and the BioSpot-VIVAS sampler, recent instruments based on bubble bursting and collection following condensational growth, respectively, attempt to minimize contact with surfaces and limit stresses on the viruses to levels occurring in the natural transmission chain. This is expected to reduce instrumental artefacts.

Motivated by the above, we use a novel chamber facility – the LAPI BREATH (Laboratory of Atmospheric Processes and their Impacts - Bioaerosol Research & Environmental Airborne Transmission Hub), together with state-of-the-art equipment for virus-laden aerosol production and collection that minimize artefacts, to study RH effects on IAV infectivity, in salt and salt-organic media. The results are compared against those obtained from droplet and bulk experiments to understand the role of virus confinement in aerosols.

## 2. Results

In this section, we first discuss the results on performance of the different means of particle generation and collection and their impact on virus inactivation, as this has implications for the interpretation of the inactivation experiments. We then discuss the results of our study on the RH dependence on IAV inactivation and compare them with literature data.

### 2.1. Impact of different nebulizers and bioaerosol samplers on virus inactivation

We evaluated the performance of nebulizers and samplers with regard to their ability to minimize virus inactivation during operation. The results are displayed in terms of IAV infectious fraction, i.e., the number of infectious viruses measured (PFU) over the total number of viruses, including those inactivated (determined by quantifying genomic copies). Comparing the results of two experiments with the same nebulizer allows comparison of performance of both samplers, and vice versa. These experiments were performed at 25% to ensure enough retention of virus infectivity.

Figure 1A shows that the BioSpot-VIVAS conserves IAV infectivity better than the BioSampler, as expected for a gentle sampling technique compared to a glass impinger. This confirms what was previously observed by Lednicky et al. (2016) for influenza viruses and by Pan et al. (2017) for bacteriophage MS2. However, in contrast to their work, our results suggest that the Collison outperforms the SLAG in terms of maintaining IAV infectivity (see Section 3 for more details).

**Figure 1.**
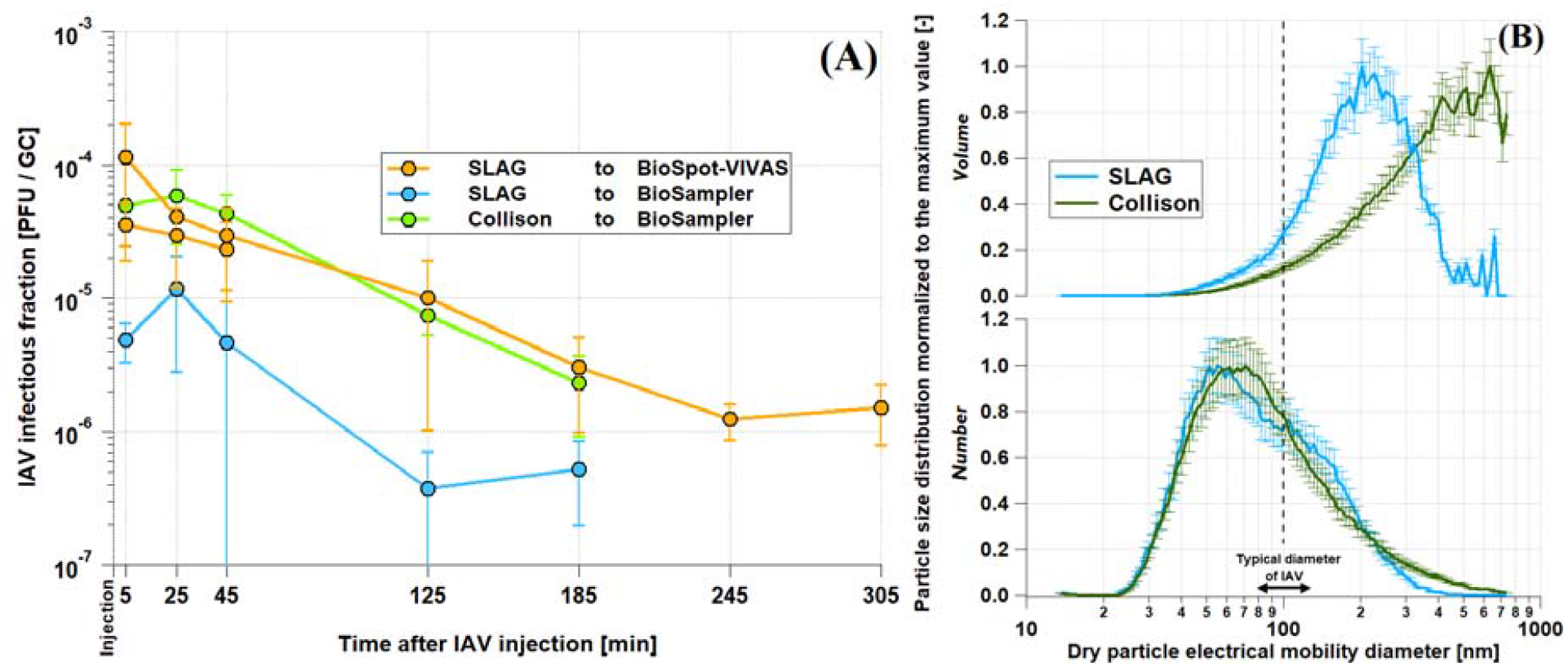
Comparison of infectivity and size distribution results from nebulizers and samplers. **(A)** IAV infectious fraction measured at 25% RH after 5, 25, 45, 125 and 185 min for different combinations of nebulizers and samplers. In order to make the data comparable, they were normalized so that the infectious fraction in the inoculated medium was identical for the three combinations. The “SLAG to BioSpot-VIVAS’’ experiment was duplicated as part of the experiments discussed in Section 2.2. The error bars indicate the 95% confidence interval on the technical triplicates. **(B)** Normalized size distributions of dry particle volume density (top panel) and number density (bottom panel) averaged over the first 5 min after virus nebulization in the chamber of the LAPI BREATH as measured by the SMPS. The normalization was performed with respect to the highest value of each size distribution. The dash line indicates the typical size of an IAV. The medium used for these experiments was PBS. Based on the results of the infectious fraction and the size distributions as well as the large amount of virus stock required to operate the Collison nebulizer, we decided to perform the experiments with the SLAG/BioSpot-VIVAS combination. The error bars indicate the counting uncertainty from the SMPS.

Dry particle size distribution measured during these experiments revealed relatively similar mode diameters between the Collison and the SLAG for number distributions (ca. 60 nm) but different volume distributions (ca. 200 nm for the SLAG and 500-600 nm for the Collison). This is owing to the different aerosol generation mechanisms between the SLAG, which emits almost no particle with a dry size larger than 400 nm, and the Collison nebulizer that generates particles of 700 nm (the maximum size measured by the SMPS) and possibly larger (Figure 1B). For both instruments, the mode of the size distribution is smaller than the nominal size of an IAV.

The results shown in Figure 1A, indicating similarities in the trend of decay between the three combinations of instruments tested, led us to the conclusion that the choice of nebulizers and samplers does not substantially influence the infectivity of viruses in the system. Although we found that the Collison outperformed the SLAG in terms of maintaining viral infectivity by about an order of magnitude, the volume of inoculated medium it requires is much higher than that of the SLAG (80 mL versus 22 mL). In a logic of optimizing experiments both in terms of protocols and results, the variations in the absolute level of IAV infectious fraction do not justify, in our opinion, the use of almost four times more virus stock. The decision to combine the SLAG and the BioSpot-VIVAS in our infectivity experiments allows us to minimize the amount of infectious material required and to realistically mimic the physiological processes of film bursting in the generation of expiratory particles (Pöhlker et al., 2023) and the growth of inhaled particles by condensation in the airways.

### 2.2. Effect of relative humidity and sucrose on airborne IAV infectivity

Infectious virus titer and total virus concentration data over the course of the 5-hour experiments performed in the LAPI BREATH are shown in Figure 2A with PBS as a medium and in Figure 2B with PBS+sucrose. IAV remained infectious up to 5 hours at low RH (10% and 25% RH), but no infectious IAV was detectable after 2 hours of exposure to higher RH levels. Total virus concentration (GC) decayed at a comparable rate among all the experiments, dictated by particle deposition on the chamber walls. Their absolute values depended on the virus stock. As described in Section 5.6, infectious titer and total virus concentration data served for the calculation of *t*_99_ (Figure 3). The highest *t*_99_ value retrieved, ca. 12 hours, corresponds to the lowest RH value (10%). At 25% RH, *t*_99_ is more than halved compared to its value at 10% RH, and drops sharply to approximately 30 min at 40% RH. A minimum of *t*_99_ ≈ 20 min is noticeable at 55% RH, followed by a modest but relatively steady increase to ca. 25 min at 70% RH, 30 min at 85% RH and 35 min at 95% RH. As shown in Figures 3 and 4, high reproducibility was achieved at each RH tested between the 5-hour experiment and the shorter 45-min duplicate (with the exception of 70% RH, where some deviation was observed). In summary, these data show a dependence of infectivity on RH with slow decay under dry conditions (*t*_99_ > 3 h at RH < 30%) and rapid decay under humid conditions (*t*_99_ < ½ h at RH > 40%), but no prominent evidence of the U-shape discussed in the literature (e.g., Tang, 2009; Yang et al., 2012).

**Figure 2.**
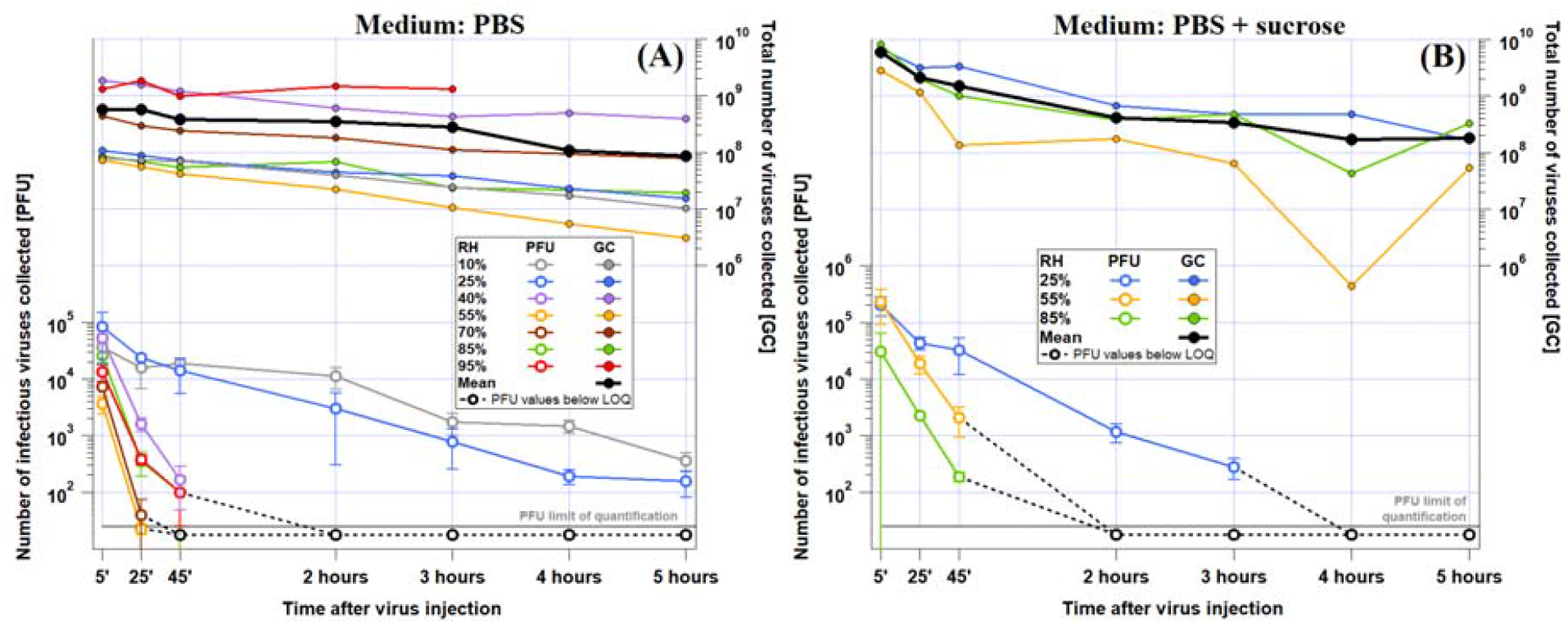
Raw RH-dependent IAV infectivity data from the LAPI BREATH. Number of infectious (PFU) and total (GC) viruses collected in Petri dishes of the BioSpot-VIVAS for **(A)** PBS and **(B)** PBS+sucrose experiments over the course of the 5 hours experiments (45-min duplicates and sucrose-supplemented medium experiments are not shown for readability). These data were used for the calculation of *t*_99_ (see Section 5.6).

**Figure 3.**
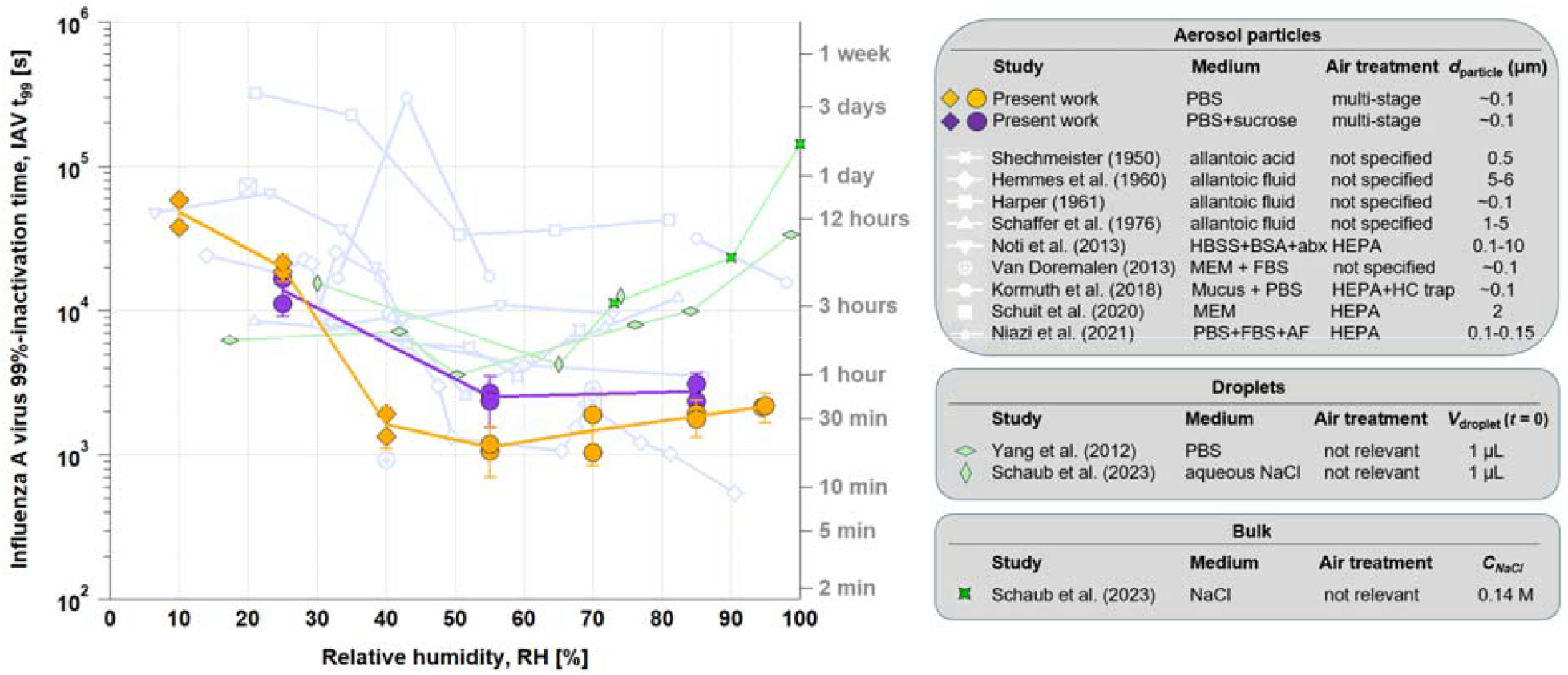
Processed RH-dependent IAV infectivity data from the LAPI BREATH in comparison to the literature. *t*_99_ versus RH in PBS (orange circles for liquid particles and diamonds for effloresced particles) and PBS supplemented with sucrose (same symbols in purple), in comparison with previous literature on RH-dependent airborne retention of IAV (light blue lines with symbols explained in the table on the right). Previous studies on RH-dependent IAV inactivation in microliter droplets (light green diamonds) and in bulk samples (dark green stars) are also depicted. The error bars indicate the 95% confidence interval on pooled experiments, combining the 45 min and the 5 hours experiments at each RH and each medium. Acronyms: *d*_particle_: particle diameter; HBSS: Hanks’ s balanced salt solution; BSA: bovine serum albumin; abx: antibiotics; FBS: fetal bovine serum; HC: hydrocarbon; AF: allantoic fluid; *V*_droplet_:droplet volume; *C*_NaCl_: sodium chloride concentration.

**Figure 4.**
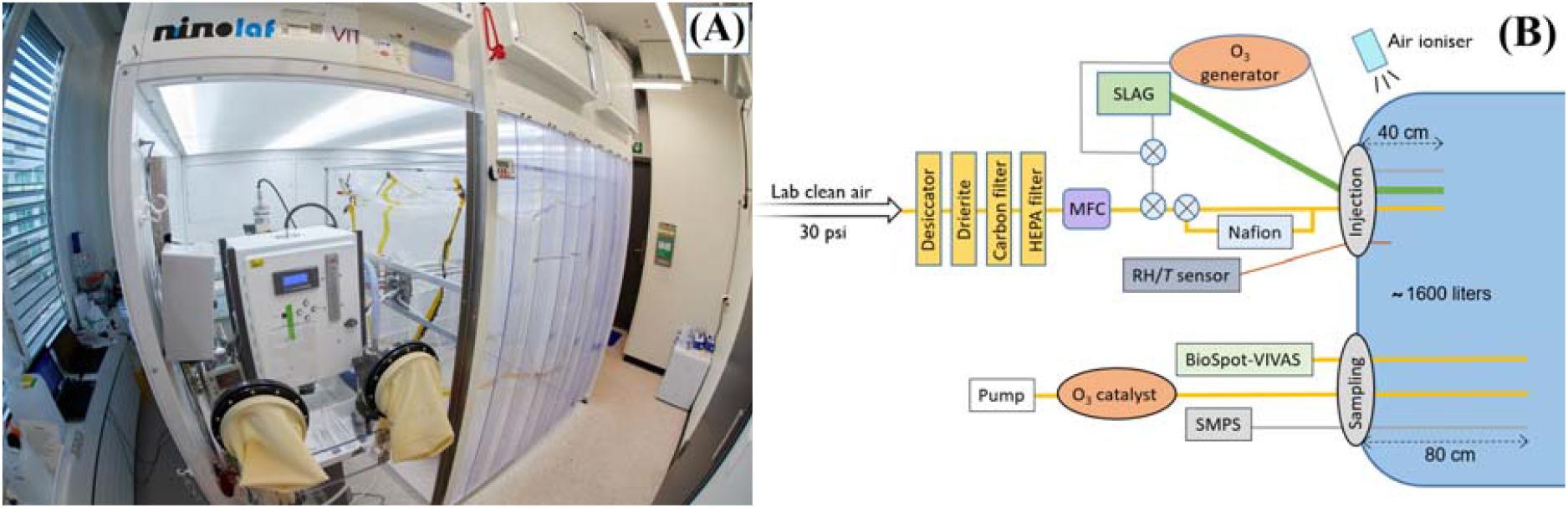
The LAPI BREATH facility and the experimental setup as used in the present work. **(A)** Photograph of the laboratory where the experiments were performed (credit: H.K. Wynn). **(B)** Schematic of the experimental setup used for airborne virus exposure experiments. HEPA filter: high-efficiency particulate air filter; MFC: mass flow controller; SLAG: Sparging liquid aerosol generator; BioSpot-VIVAS: viable virus aerosol sampler; SMPS: scanning mobility particle sizer; RH/*T*: relative humidity/air temperature sensor. The crossed circles represent manual three-way valves.

Figure 3 compares *t*_99_ of our aerosol experiments with the results of studies using microliter droplets (green symbols) by Yang et al. (2012) and Schaub et al. (2023). These studies also investigated the infectivity of IAV in aqueous solutions, namely PBS and aqueous NaCl solution, respectively. They show good agreement with each other, although the first study assumes first-order kinetics both before and after efflorescence, while the second is based on measured kinetics during the equilibrium phase after efflorescence only. Both studies agree well with each other and come to a similar interpretation, namely that in a saline droplet the increased salt concentration resulting from evaporation is an inactivating agent for the viruses, which is attenuated by efflorescence, so that minimal infectivity is to be expected at a water activity *a*_*w*_ just above the efflorescence point, when water is still present and dissolved salt concentrations are at their highest. This is around *a*_*w*_ ∼ 0.5 - 0.6, corresponding to ERH ∼ 50% - 60% under equilibrium conditions. The agreement between the experiments with aerosol particles and deposited droplets is reasonably good at RH < 30%. However, at medium and high humidity, the *t*_99_ derived from the droplet measurements is consistently up to an order of magnitude higher than *t*_99_ in the aerosol measurements.

Bulk experiments, with *a*_*w*_ equal or very close to 1, are consistent with large droplet experiments at 100% RH given the very dilute nature of the solution and the large salt-to-virus mass ratio found in both types of experiments. The *t*_99_ value reported by Schaub et al. (2023) in bulk reaches more than 10^5^ s, higher than any other value retrieved at lower RH. To compare to the results of the LAPI BREATH aerosol experiments, where it is technically impossible to achieve 100% RH conditions in the chamber, we performed additional bulk experiments corresponding to 90% and 73% RH, which yield *t*_99_ a factor of 6 and 13 lower than at 100% RH, respectively, but still about an order of magnitude slower than the aerosol in the chamber.

Using a purely saline solution medium such as PBS helps reduce experimental complexity by limiting the number of compounds that can influence IAV inactivation kinetics in the matrix and by buffering against pH changes. However, many other substances are present in physiological aerosol matrices, such as airway lining fluid or saliva. We tested the potential impact of sucrose on the infectivity-RH relationship by adding it to the same initial PBS medium (prior to aerosolization). Sucrose was chosen as a representative of organic substances, which are known to be abundant in respiratory tract lining fluid. The concentrations of NaCl and sucrose in the inoculated medium examined were 8 g/L and 64 g/L respectively, corresponding to an NaCl:sucrose dry mass ratio of 1:8. This NaCl:sucrose ratio is sufficient to suppress efflorescence at 55% RH (Schaub et al., 2023).

The experiments in which we supplemented the PBS medium with sucrose were performed at 25%, 55% and 85% RH. Figure 3 shows that the effect of sucrose is small, and, if present, there is a protective effect on IAV at intermediate RH, increasing *t*_99_ slightly from approximately 20 min to 40-45 min, but with no observable impact at 25% and 85% RH. The effect of sucrose supplementation can thus be described as a further “flattening” of the U-shape.

In EDB experiments using PBS, efflorescence occurred at 43.7±1.5% RH in droplets with a radius of 8 µm just prior to efflorescence. For comparison, a particle composed of aqueous NaCl with an initial radius of 6.4 µm effloresced at

44.3±1.5% RH (Luo et al., 2022). We conclude that during the LAPI BREATH experiments, the PBS particles were in the metastable liquid state at RH above 50%, whereas efflorescence most likely occurred during the experiments performed at 40% RH or below. The increase in matrix viscosity caused by the high proportion of sucrose added to the medium directly impacts efflorescence. Similar to other organic/inorganic mixtures (Song et al., 2013), we consider it very likely that aerosol particles originating from a PBS+sucrose medium in the LAPI BREATH only effloresced at very low RH or did not effloresce at all, regardless of RH. However, we cannot exclude the formation of microcrystals, as described by Schaub et al. (2023), which do not grow because of the high viscosity and low diffusivity.

## 3. Discussion

### 3.1. Performance of nebulizers

The results presented in Section 2.1 indicate increased infectivity when nebulizing IAV in saline medium with the Collison nebulizer compared to the SLAG. This was unexpected, given the expectations based on the two aerosolization methods - strong shear forces and high intensity shocks with impaction versus bubble bursting with sparging liquid. This also contradicts previous work from Paton et al. (2022), who reported a higher viable fraction of SARS-CoV-2 virus for the SLAG compared to the Collison nebulizer. Particle size distribution from both instruments (Figure 1B) may explain this counterintuitive result. Zuo et al. (2013), who investigated avian influenza virus size distribution in aerosol particles emitted by a Collison nebulizer, showed that both infectious and total virus size distributions follow the aerosol volume size distribution. Interestingly, they also showed that the infectious fraction in aerosol particles normalized to the infectious fraction in the inoculated medium, was higher over the dry particle diameter range between 300 and 450 nm than between 100 and 300 nm. As the SLAG produces a much lower mass fraction of 300 to 400 nm aerosol particles compared to the Collison nebulizer, it is possible that the overall infectious fraction observed from the SLAG appears lower than that of the Collison nebulizer.

Performance comparison studies between the SLAG and the Collison, similar to the one reported in Figure 1A, favored the SLAG for nebulizing bacteria (Rule et al., 2009 with *Pantoea agglomerans*; Zhen et al., 2014 with *Escherichia coli*). This is most probably due to the strong inertia of the bacteria tested, whose size is about an order of magnitude larger than IAV, thus potentially increasing the damage caused by impaction forces compared to viruses in the Collison nebulizer. It is however also possible that, although the very small mass of IAV with limited inertia avoids structural damages within the high-velocity flows of the Collison nebulizer, other forces may cause damage. One possibility is that surface tension effects from viruses contained within evaporating droplets may disrupt enveloped viruses and inactivate them (Coleman et al., 2024). The same study concludes that osmotic pressure from dissolved salts still may dominate inactivation of viruses, not because of differences in their magnitude, but rather the exposure time to each force. However, given that the SLAG generates smaller droplets than the Collison nebulizer, owing to the film-bursting mechanism, the viruses may reside near the drop-air interface for much longer periods of time (more partially engulfed viruses compared to more fully immersed viruses produced by the Collison nebulizer), hence be exposed to damaging surface tension for longer. Shear stress does not seem to play a significant role, as it has a much smaller magnitude than surface tension and osmotic forces. These hypotheses suggest that it is not only the softness of the nebulization, but also the particle size affecting the decay during the particle production; they are consistent with the large differences in infectivity between the Collison and the SLAG when nebulizing bacteria (Rule et al., 2009; Zhen et al., 2014).

### 3.2. Disagreement between aerosol particle and microliter droplet experiments

In Figure 3, we compared microliter droplet experiment results from Yang et al. (2012) and Schaub et al. (2023) with our results from aerosol particle experiments, all using saline media. The fast equilibration of aerosol particles and microliter droplets to ambient RH above ∼50 % should, in principle, make the results of both types of experiments comparable. Furthermore, at such RH, efflorescence cannot occur, thus avoiding the additional complexity existing at low RH. We can only speculate about the reasons for the discrepancy of up to an order of magnitude between *t*_99_ from microliter droplet and aerosol particle experiments. Reasons could be the dependence of *t*_99_ on (i) the initial virus titer leading to a protective effect, (ii) the pH of PBS, which in turn is a function of *a*_*w*_ and thus of RH, and (iii) the location of the viruses on the surface of the very small aerosol particles exposing them to the air and to strong tension forces not present in the larger droplets or bulk. Each of these hypothetical reasons is discussed in more detail in the Supporting Information, and it is shown that in combination they can indeed bring the measurements into agreement (see Figure S2). Further work is needed to assess the validity of these hypotheses. In summary, bulk or microliter droplet experiments cannot currently be reliably extrapolated to study the behavior of airborne IAV with respect to inactivation kinetics. Conversely, aerosol chamber experiments rely on the choice of particle size, which may influence their results.

### 3.3. IAV infectivity versus RH relationship and metrics for respiratory disease transmission

The remarkable impact of RH on the stability of IAV infectivity in a saline matrix is clearly shown by the decrease of *t*_99_ by 1.5 orders of magnitude during the transition from dry to humid conditions (see Section 2.2). As shown in Figure 3, several studies reported a similar RH-dependence, although to a lesser extent (Shechmeister, 1950; Harper, 1961; Hemmes et al., 1962; Schaffer et al., 1976; Noti et al., 2013). It should be noted, however, that important experimental parameters were sometimes poorly described or even unknown in previous work, such as the exact composition of the medium or of the applied ambient air, making precise comparisons between these studies difficult.

The change in IAV infectivity between intermediate and high RH has been widely discussed in a number of studies. To describe the results from Schaffer et al. (1976), Tang (2009) used the term “asymmetrical V-shaped curve”, giving rise to a long-standing debate over whether such a V-shaped or U-shaped relationship between IAV infectivity and RH actually exists. It is important to note that in these previous reports, infectivity was not expressed as *t*_99_, but as a normalized titer *C*(*t*)/*C*(0) measured at a certain time *t* (typically 1 h). Due to the exponential dependence *C*(*t*) = *C*(0) exp(-*t*/*t*_99_ ln(100)), a shallow minimum in *t*_99_ can result in a pronounced minimum in *C*(*t*) for long times *t. t*_99_ however is a preferred metric for the quantification of infectivity, as it is derived from the measurement of the time course with several temporal support points, hence less prone to uncertainty than a specific time instant that corresponds to titer measurements. A decrease by two orders of magnitude indicated by *t*_99_ represents systematic and substantial decrease in magnitude relevant to health policy.

The unit chosen by Schaffer et al. (1976) to describe the conservation of infectivity was the “% recovery”, i.e., *C*(*t*)/*C*(0) expressed as a percentage of IAV still infectious after a certain duration of exposure in the chamber. By converting units of retention of IAV infectivity to *t*_99_ from the data of Schaffer et al. (1976) as well as several other studies, we observed that V- or U-shapes largely disappear (Figure 3) and lead to a very slight increase in IAV infectivity with RH increasing from intermediate to high values - or even to no increase at all. Conversely, we calculated inactivation rates after 1 hour and 3 hours of exposure based on our data from Figure 2A, 2B and show the results in Figure S1. This leads to clear U-shaped curves. The visual impression of a V- or U-shape therefore depends on whether one specifies *t*_99_ (and thus the logarithmic quantity) or C(t)/C(0), which magnifies the RH dependency exponentially with time. The normalized titer C(t)/C(0), although widely used and mathematically equivalent to specifying *t*_99_, is a less suited measure to link the maintenance of viral infectivity to the risk of indoor respiratory disease transmission because it depends on the duration of exposure, which complicates its interpretation. Instead, *t*_99_ can be directly related to the suspension time of aerosol particles in indoor transmission scenarios, as it gives an indication of the time after which the risk of transmission is substantially reduced (by two orders of magnitude), without any dependence on exposure duration.

The mechanistic underpinning of the influence of RH on IAV infectivity has only recently begun to be uncovered (Yang and Marr, 2012; Morris et al., 2021). In saline suspensions, where the degree of complexity is minimal, recent studies have suggested that the protection of IAV at low RH is not afforded by the salt crystal itself, but rather by the effect of the salt crystal depleting the salt from the surrounding solution (more precisely the NaCl molality) in which the virus is immersed (Niazi et al., 2021; Schaub et al., 2023). The increase in salinity in the particle leads to strong inactivation at intermediate RH (55% to 70%), while the reduced loss of water vapor from the exhaled particles at very high RH (85% to 95%) keeps salt concentrations sufficiently low and thereby *t*_99_ high.

### 3.4. Impact of organics on the retention of IAV infectivity

The addition of sucrose to our PBS medium led to a further flattening of the already only slightly concave *t*_99_ curve as a function of RH (purple and orange curves in Figure 3). Several recent studies reported very weak and RH-independent IAV inactivation when supplementing their medium with proteins (Kormuth et al., 2018; Schuit et al., 2020; Dubuis et al., 2021). The present study supports that organic species in the aerosol tend to protect IAV at intermediate RH, albeit not to a large extent, thereby reducing the effects of RH on IAV inactivation. Note that we decided not to extract any *t*_99_ value from the study of Dubuis et al. (2021) due to the large dispersion in the data and the very low inactivation rates leading to *t*_99_ values which can be considered infinite.

As the properties of the aerosol particles in Figure 3 show, all the previous studies used organic-rich matrices, mostly by addition of proteins. Only two studies, Shechmeister (1950) and Schaffer et al. (1976), used a null or low proportion of proteins relative to salts. However, the medium they used contained large amounts of other organics, which may explain why the RH-dependence derived from their measurements is stronger than what we observed with PBS, but close to PBS + sucrose.

### 3.5. Recommendations for future studies

The fact that the *t*_99_-RH relationship from our aerosol particle experiments resulted in the lowermost curve in Figure 3 compared to literature data (except for a few data points from Hemmes et al. (1962) at high RH) is an indication that we succeeded in constructing an experimental setup that excludes any potential protective agent to IAV from the matrix (condensed phase) or the air (gas phase). This gives us the ability to individually characterize single protective or inactivation factors by modifying the experimental conditions one by one and, ultimately, it gives us confidence that the LAPI BREATH will enable a realistic simulation of virus-containing aerosol in relevant matrices, thus substantially contributing to the understanding of the factors that control airborne transmission of viruses in an indoor environment. Drawing on the experience gained in the design, operation and optimisation of this setup, we provide the following recommendations for the experimental study of the airborne transmission of viruses:

- The large volume, and in particular the large vertical dimension of the LAPI BREATH makes it possible to avoid the use of an aerosol resuspension system. Although such dimensions require large biosafety cabinets, they allow the natural particle size-dependent sedimentation rate to be left unchanged, which is not the case with rotating drums, fan-equipped chambers and single-particle levitation techniques. Static chamber experiments are therefore the only type that allow distribution-averaged infectivity results to be obtained. The only strong difference we notice between an experiment in the LAPI BREATH compared to an indoor environment is the very high concentration of particles (and virus-containing particles) aloft in our system. Although the effect of particle concentration on virus inactivation has not been studied to our knowledge, we did not observe evidence of substantial coagulation in the size distribution measurements from the SMPS and therefore assume that interactions between IAV-containing particles did not alter infectivity results.
- The experiments to study the retention of IAV infectivity in suspended or deposited droplets are not able to accurately mimic specific conditions found in aerosols (Haddrell and Thomas, 2017) and any conclusion on the conservation of infectivity of airborne viruses based on their results should therefore be taken with caution.
- We advocate the use of samplers that avoid contacts of airborne infectious pathogens with surfaces, especially if these contacts are associated with high speeds/pressures in order to avoid structural damages responsible for loss of infectivity.
- In order to obtain infectivity results that consider the entire kinetics of each experiment and make them independent of exposure time, we also recommend taking samples at different times during an experiment. Along the same lines, we believe that infectivity results should be expressed at *t*_99_, thus providing a relevant measure for the indoor suspension time of respiratory viruses.
- Following recommendations from Santarpia et al. (2020) and Groth et al. (2024), medium and air composition used during the experiments should be described as best as possible, in particular the presence of proteins and organic matter in the medium and trace gases in the air (controlled and/or measured concentrations, or failing that, a description of the filtration system used), as they have an influence on the retention of IAV infectivity.

## 4. Conclusions

Despite decades of research, there remains great difficulty in identifying and understanding the factors that control the spread of influenza epidemics, which occur each year in both hemispheres. In this study, we aimed to simulate exhalation, airborne residence, and re-inhalation of IAV-containing particles as accurately as possible. As an initial step, we specifically used purely saline aerosol particles to avoid the complexity of natural matrices. We revealed large discrepancies between aerosol-borne viruses and previous microliter droplet experiments performed with similar matrices. The discrepancies between aerosol and deposited droplet experiments need to be explained in order to obtain a reliable model for aerosol transmission. We confirmed literature findings showing that intermediate and high RH levels (above ca. 40% RH) are significantly more favorable for inactivating airborne IAV in saline matrices than dry conditions (< 30% RH). In a second step, we have quantified the protective effect of sucrose in the medium, confirming previous studies showing the delay of IAV inactivation by organic material, especially proteins, and thus mitigating the effect of relative humidity on IAV infectivity. Further work with matrices representative of the human respiratory system will reveal whether maintaining indoor environments at a specific RH may be an effective mitigation strategy to reduce the risk of airborne transmission of IAV. We also introduce the LAPI BREATH facility, an experimental setup that allows us to mimic the airborne transmission of IAV using new generation instrumentation. We tested it in comparison to standard techniques and showed that this setup is effective for carrying out experiments towards disentangling the impact of individual factors in the inactivation of aerosol-borne IAV. Finally, we put our results into the context of literature data and showed that a U-shape relationship as discussed in the past largely disappears when displaying results in terms of *t*_99_, which we believe is the most relevant metric regarding the risk of virus transmission. The gaps in our understanding of airborne virus transmission require transdisciplinary efforts in terms of experimentation and modelling, where the LAPI BREATH can make an important contribution.

## 5. Experimental Procedures

### 5.1. Virus preparation

Experiments were conducted with IAV strain A/WSN/33 (H1N1 subtype). Madin-Darby Canine Kidney (MDCK) cells (ThermoFisher Co, Waltham, MA, USA) maintained in Dulbecco’s modified Eagle’s medium (DMEM, Gibco, ThermoFisher) supplemented with 10% Foetal Bovine Serum (FBS; Gibco), and 1% Penicillin-Streptomycin 10,000 U/mL (P/S; Gibco) were used for virus propagation. Confluent MDCK monolayers were washed and inoculated with IAV at a multiplicity of infection of 0.001 for 48 to 72 h in OptiMEM (Gibco) supplemented with 1% P/S and 1 μg/mL TPCK trypsin (T1426, Sigma-Aldrich Inc., Saint-Louis, MO, USA). The culture supernatant from the infected cells was cleared by centrifugation (2,500 ×g, 10 min). Subsequently, the virus was concentrated and purified by pelleting through a 30% sucrose cushion at 112,400 ×g in a SW31Ti rotor (Beckman) for 90 min at 4 °C. The resulting pellets were soaked overnight at 4 °C in phosphate-buffered saline (PBS, ThermoFisher, 18912014) and then fully resuspended by pipetting up and down. Concentrated IAV stocks were quantified using plaque assay (described below), resulting in a titer of ∼10^11^ plaque forming units per milliliter (PFU/mL). The virus was aliquoted and frozen at -80 °C for long-term storage.

### 5.2. Medium preparation and sample handling

Two different media were used in this study: PBS and PBS supplemented with sucrose (see the exact composition of the media in Table S1). The sucrose solution was prepared by dissolving 64 mg/mL of sucrose (ThermoFisher BioReagents, BP220-1) in PBS (i.e., the sucrose to NaCl ratio is 64 mg/8 mg ≈ 4 mol/3 mol). A volume of 22 mL of medium was then placed in a 100-mL screw-capped container and 110 μL or 220 μL of virus stock was spiked in the medium (in order to reach a titer of ∼10^9^ PFU/ml). The virus suspension was kept at 4 °C during preparation and transportation to the LAPI BREATH. Hereinafter, we refer to the prepared solution as “medium” before virus inoculation and “inoculated medium” after the virus stock is spiked. We use the term “matrix” to refer to the liquid in dynamic physico-chemical evolution contained in the aerosol particles, which differ from the inoculated medium due to the evaporation process.

To ensure that the initial virus concentration was comparable between experiments, 25 μL of the inoculated medium was sampled into 2.5 mL of PBSi (PBS for infection; PBS supplemented with 1% P/S, 0.02 mM Mg^2+^, 0.01 mM Ca^2+^, and 0.3% bovine serum albumin (BSA, Sigma-Aldrich, A1595), with a final pH of ∼7.3) and kept at 4 °C in a 50-mL tube for the duration of the experiment. This allowed us to take account of any natural decay of the virus in bulk solution. After aerosolization (or nebulization; see Section 5.3) of the inoculated medium into the LAPI BREATH, individual samples were collected across a 5-hour time course. Each sample was recovered in a Petri dish (35_⍰_10 mm Petrischale, catalogue number D210-15, Biosystems Switzerland AG, Muttenz, BL, Switzerland) filled with 2.5 mL of PBSi, transferred in a 50-mL centrifuge tube and stored on ice until the end of the experiment. The samples were then divided into 3 portions: 3_⍰_200 μL for infectivity quantification by plaque assay (protocol described in Section 5.4), 200 μL for total virus quantification by RT-qPCR (reverse transcription-quantitative polymerase chain reaction; protocol described in Section 5.5) to assess the wall losses, and the remaining volume (∼1.7 ml) for backup storage. All the sample aliquots were frozen at -20 °C until use.

### 5.3. Aerosol particle experiments: aerosolization, exposure and sampling

The core of the LAPI BREATH is a ca. 1.6 m^3^ (length/width/height = 1.36 m / 0.82 m / 1.44 m) polytetrafluoroethylene (PTFE) chamber suspended from a metal frame inside a biosafety enclosure (Figure 4A), based on established knowledge from the aerosol chamber community (e.g., Kaltsonoudis et al., 2019; Doussin et al., 2023). Leakage out of the chamber was minimized by plugging leak points with vacuum sealed strips – identified by slightly overinflating the chamber using a mixture of air and helium, and detecting the leaks with a portable helium leak detector. Prior to each experiment, the chamber was flushed with dry purified air. The laboratory clean air inlet provided outdoor air taken from the building roof (fourth floor), cooled or heated to 21°C and filtered using a classical particle filter (class F9). To avoid interaction with uncontrolled trace gases in the chamber air, such as acids (Luo et al., 2022), the laboratory clean air was additionally dried using drierite and filtered with activated carbon and HEPA filters (Figure 4B). Experiments commenced after cleanroom conditions with number concentrations below 3 cm^-3^ were reached for particles > 7 nm. The introduction of virus-containing aerosol particles into the chamber was conducted with the chamber being slightly deflated (by pumping out ∼100 liters of air) to prevent leakage of infectious viruses from the chamber. The chamber temperature was maintained between 23 and 25 °C by keeping the laboratory temperature constant during the experiments. RH was controlled by injecting water vapor through a Nafion humidifier (model FC100-80-6MKS, Permapure LLC, Lakewood, NJ, USA). Air temperature and RH were monitored using a probe (model SP-004-2, Omega Engineering, Norwalk, CT, USA) with a manufacturer-given accuracy of ±0.3 °C for temperature, and ±2.5% RH (±3.5% RH) when lower (greater) than 80% RH.

The continuous downward airflow inside the biosafety enclosure leads to electrostatic charging of the chamber walls, which promotes the loss of aerosol particles. To neutralize these charges, an air ionizer (model SL-001, Dr. Schneider Holding GmbH, Kronach-Neuses, Bavaria, Germany), which uses discharges from a high-frequency piezoelectric transformer to ionize a stream of air, continuously blew air down the PTFE foil outside the chamber during experiments.

Virus-containing aerosol particles were introduced in the chamber of the LAPI BREATH using a 90 mm sparging liquid aerosol generator (SLAG, CH Technologies Inc., Westwood, NJ, USA), which forms an aerosol from a suspension liquid by bursting bubbles. This mimics the natural formation of aerosols by air-liquid interface systems, such as the respiratory tract, natural and man-made bodies of water etc. (Alsved et al., 2020; Pöhlker et al., 2023). A 22 mL volume of viral suspension was placed on the frit plate, through which purified air was sent with a flow rate of 30 L/min. The SLAG was connected to the chamber by a tube of 20-mm internal diameter and the point of injection was located 40 cm from the chamber wall, so that the concentration of aerosol particles injected towards the center of the chamber homogenized as quickly as possible.

A pulsed injection was performed with a short nebulization time, namely 30 seconds with PBS and 1 min when sucrose was added to PBS. The aerosolization time was adjusted because the presence of sucrose in the inoculated medium generates less aerosol particles per second compared to PBS (likely due to modifications of the viscosity and the surface tension of the inoculated medium). The short time span between the first and last virus injected into the chamber in relation to the sampling time ensures that all viruses in a collected sample have been exposed to similar conditions.

During aerosol exposure, air was sampled 80 cm away from the chamber wall, as close to its center as possible, using stainless steel tubes inside the chamber and silicone conductive tubings between the chamber and the instruments. To minimize particle loss, the inner diameter of tubes and tubings was 0.635 cm (¼ inch) for instruments with a flow rate lower than ca. 4 L/min and 0.95 cm (⅜ inch) for higher flow rates.

The particle size distribution was measured continuously using a scanning mobility particle sizer (SMPS), which consists of a differential mobility analyser (DMA model TSI long, TSI Inc., Shoreview, MN, USA) and a condensation particle counter (CPC model 3772, TSI Inc., Shoreview, MN, USA). The DMA scanned the particle mobility diameter between 13.6 and 736.5 nm, forming a monodisperse aerosol flow whose concentration is measured by the CPC.

Virus sampling was performed using a viable virus aerosol sampler (BioSpot-VIVAS model BSS310, Aerosol Devices Inc., Fort Collins, CO, USA; see Lednicky et al., 2016), which, before sampling, grows aerosol particles from as little as a few nm to micron-sized droplets via water vapor condensation. This ensures that the particles are collected gently on a Petri dish filled with 2.5 mL of PBSi. The supersaturation within the instrument column is produced by the injection of liquid water and the heating or cooling of different stages: the conditioner, initiator, moderator, nozzle, and sample were respectively set to 5, 45, 13, 25 and 13 °C. Samples were collected with a flow rate of 8 L/min. To ensure efficient condensation of water vapor onto virus-containing aerosol particles in the BioSpot-VIVAS column during sampling, the aerosol concentration was kept below 50000 cm^-3^ in the aerosol chamber. A higher particle concentration could cause excessive competition for water vapor within the column, which would lead to a degrading collection efficiency. After collection, the samples were poured into conical 50-mL centrifuge tubes and placed in an ice bucket until the end of the experiment, and subsequently aliquoted and frozen at -20 °C. After each experiment, the chamber was decontaminated for one hour using a combination of UV radiation and flushing with ozone-rich air (6 ppm for 1h) produced by an ozone generator. Sampling of decontaminated air from the chamber post-experiment confirmed this process was sufficient to remove any infectious viruses. The ozone-enriched air containing the inactivated viruses was then passed through an ozone catalyst and replaced by purified air to prepare for the next experiment.

We measured the retention of IAV infectivity in aerosol at seven RH levels between 10% and 95%, keeping all parameters and experimental conditions constant and identical except for RH. For each RH, an experiment was conducted over 5 hours, with air samples taken from the chamber after 0, 20, 40, 120, 180, 240 and 300 minutes of exposure. For each time, air was drawn from the chamber into the BioSpot-VIVAS for 10 consecutive minutes. A shorter duplicate experiment was also performed for each RH and stopped after the 40-minute sampling time point. These shorter replicates were sufficient to calculate independent inactivation rates at each of the RH settings, with the exception of 55% RH, where only one of the two experiments yielded infectivity results above the limit of quantification (LOQ) after 25 min of exposure.

We compared the loss of infectivity caused by a 6-jet Collison nebulizer versus a 90 mm SLAG during aerosolization, and by the BioSampler versus the BioSpot-VIVAS during sampling (Section 2.1). Three experiments were performed by permuting the pairs of nebulizer and sampler. However, we did not compare the two samplers by running them simultaneously: the BioSampler was utilized with a flow rate of 12.5 L/min, which in combination with the 8 L/min of the BioSpot-VIVAS was too high to allow efficient sampling. The three experiments were therefore carried out separately, but under identical conditions. Due to the large amount of virus stock required to operate the Collison nebulizer, only one experiment using this instrument was performed.

### 5.4. Titration of infectious viruses in the aerosol particles and droplets

Determination of the infectious virus concentration in each sample obtained from the aerosol particle and the microliter droplet experiments was done using a plaque assay with MDCK cells as described previously (David et al., 2023). Briefly, 12-well plates containing cellular monolayers were washed with PBS and infected with IAV samples. Prior to infection, the samples were serially diluted in PBSi (see Section 5.2). A negative control was performed on an additional plate, using PBSi as a blank sample. Infected monolayers were incubated for 1h at 37 °C with 5% of CO_2_, and manually agitated every 10 min. The medium containing non-attached viruses was removed and the cells were covered by an agar overlay (modified Eagle’s medium, MEM, supplemented with 0.5 μg/mL of TPCK-trypsin and 2% of Oxoid agar). Plates were incubated for 72 h at 37 °C with 5% CO_2_ and cells were then fixed for 20 min using PBS supplemented with 10% formaldehyde (Sigma-Aldrich, 47608-1L-F) and stained for 10-20 min with a 0.2% crystal violet solution (Sigma-Aldrich, HT901-8FOZ) in water + 10% methanol (ThermoFisher Chemical, M-4000-15) to allow for plaque enumeration. The LOQ for plaque assays was 10 PFU/mL. Non-Detectable values were treated according to the recommendations of Hornung and Reed (1990). In the case where only one of the three PFU triplicates was below LOQ, the value 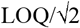 was assigned to it. If only one of the three triplicates was above the LOQ, its value was disregarded and considered below the LOQ.

### 5.5. Quantification of physical losses using RT-qPCR

RNA extractions from 140 μL of samples were performed, using the QIAamp Viral RNA Mini extraction kit (52906, QIAGEN GmbH, Hilden, NW, Germany) according to manufacturer’s instructions. Extracted nucleic acids were eluted in 80 μL of elution buffer, and stored at -20 °C until analysis. An additional negative extraction control was always performed to ensure the cleanliness of the solutions. Reverse-transcription and amplification were performed using the One Step PrimeScript™ RT-PCR Kit (RR064A, Takara Bio Inc., Kusatsu, JP-25, Japan) with the following forward primer (5’-ATGAGYCTTYTAACCGAGGTCGAAACG-3’) and reverse primer (5’-TGGACAAANCGTCTACGCTGCAG-3’) that target a 244-base amplicon of the IAV M segment.

The RT-qPCR mix was composed of 7.5 μL of 2x One-Step SYBR RT-PCR Buffer, 0.3 μL of Takara Ex Taq HS (5 U/μL stock), 0.3 μL PrimeScript RT enzyme mix, 0.3 μL forward and reverse primers (10 µM stocks), and 3.3 μ l RNase-free water, to which 3 μL of extracted RNA sample was added. The RT-qPCR was performed in a Mic Real-Time PCR System (Bio Molecular Systems, Upper Coomera, QLD, Australia) with the following parameters: 2 min at 50 °C, 10 min at 95 °C, 15 sec at 95 °C followed by 60 sec at 60 °C for 40 cycles. A final dissociation step from 55 °C to 95 °C at 0.3 °C/s was performed for the melting curve analysis. A Gblock gene fragment (Gblock SA, Ath, WHT, Belgium), as described in Olive et al. (2022), was used to create a standard curve for quantification over a range of 10 to 10^7^ genomic copies per microliter (GC/μl). The absence of inhibition was regularly checked using serial dilutions of the samples. Pooled standard curves were analysed using the Generic qPCR limit of detection (LOD) / limit of quantification (LOQ) calculator (Klymus et al., 2020). The average slope of the standard curve was -3.40 and the average intercept was 33.46. R^2^ of individual standard curves were typically ≥ 0.99, with PCR efficiencies from 86 to 95%. The LOQ was determined at 10 copies/reaction. The LOQ is defined as the lowest standard concentration with a coefficient of variation smaller 35%. All RT-qPCR procedures followed MIQE guidelines (Bustin et al., 2009; see Table S2).

### 5.6. Data treatment

To determine the inactivation kinetics of IAV at each RH, the number of infectious virus (PFU) and total virus (GC) collected in Petri dishes of the BioSpot-VIVAS were determined at all time-points. As we will show below, the inactivation can be approximated by a first-order loss process. The first-order inactivation rate constant for infectious virus, *k*_*PFU*,_ was determined from least square fits to the log-linear portion of the inactivation curves:

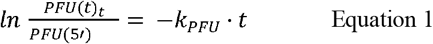

Here, *PFU*(t) is the infectious virus titer, i.e. the number of infectious viruses collected in the Petri dish of the BioSPot VIVAS, at time t (t is assigned as the median time of the 10 min sampling for each time-point), *PFU*(5’) is the initial virus titer from the first sampled time-point (median of 0 – 10 min post-aerosolization) and *k*_PFU_ is the inactivation rate constant. As samples were taken over the course of 5 hours, some physical losses of virus-containing aerosols (due to gravitational settling and/or deposition on the chamber walls) are to be expected. As IAV genome copies are not degraded in aerosols over our measured time-course, GC is only reduced by physical removal of virus-containing aerosols from the sampled air. Thus, the first-order physical loss rate constant, *k*_*GC*_, is:

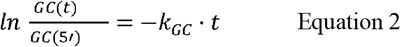

Here, *GC*(t) is the concentration of genome copies, i.e. the total number of viruses collected in the Petri dish of the BioSPot VIVAS, at time t, *GC*(5’) is the initial genome concentration from the first sampled time-point (*t*_*S* ′_) and *k*_*GC*_ is the physical loss rate constant. Note that we determined a single *k*_*GC*_ from all the experiments with each type of medium, since we did not observe any influence of RH on *k*_*GC*_ but different particle size distributions observed with pure PBS versus PBS+sucrose as a medium could lead to different decay in GC.

Lastly, the corrected inactivation rate constant, *k* _*PFU-GC*_ was calculated as follows:

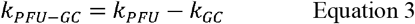

IAV infectivity results are then reported as the 99% inactivation time:

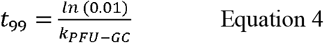

This is the time required to reduce the titer by two orders of magnitude, which we consider to be the most relevant metric (see Section 3.3).

### 5.7. Determination of the phase of particles during aerosol chamber experiments in the LAPI BREATH

To estimate whether the aerosol particles are in the liquid or crystalline state under the conditions in the chamber of the LAPI BREATH, we performed auxiliary experiments with larger particles (a few micrometers) in an electrodynamic balance (EDB). The EDB has been described by Luo et al. (2022) and in more detail by Steimer et al. (2015). To obtain an upper limit for the efflorescence relative humidity (ERH), it is sufficient to show in experiments with a larger particle that at an experimentally determined RH, the particle exists in a metastable (supersaturated) liquid state: smaller particles will always effloresce homogeneously at the same or lower RH. We performed two experiments during which Mie resonance spectroscopy was used to continuously size the particle. The first experiment consisted in levitating a single particle of undiluted PBS in the EDB and reducing RH from above 80 % down to 30 % over a time period of 10 hours at a constant temperature of 291 K. This slow drying ensures equilibrium between particle and gas phase. The micron-sized particles effloresced between 45% and 50% RH. Hence, we conclude that the submicron particles in the aerosol chamber effloresce below 45% RH.

We also performed several experiments with added sucrose at various dry mass ratios of sucrose to NaCl in the particles using the same temperature and rate of drying. No efflorescence was observed when the mass ratio exceeded 3:1. As the measurements with sucrose in the present work had sucrose:NaCl = 8:1, no efflorescence is expected.

