## Supplemental Tables, Text and Figures for "Dependence of aerosol-borne influenza A virus infectivity on relative humidity and aerosol composition"

Table S1. Composition of the media used in the present study.

| **Medium** | **Component** | **Concentration [g/L]** |
| --- | --- | --- |
| Phosphate-buffered saline (PBS)  Product number 18912 from ThermoFisher/Gibco | Sodium chloride | 8.120 |
|  | Phosphate (as sodium phosphates) | 0.950 |
|  | Potassium chloride | 0.201 |
| PBS + sucrose | Sucrose | 64.0 |
|  | Sodium chloride | 8.120 |
|  | Phosphate (as sodium phosphates | 0.950 |
|  | Potassium chloride | 0.201 |

Table S2. Check-list of experimental details as requested by MIQE guidelines according to Bustin et al. (2009).

| ITEM TO CHECK | PROVIDED  Y/N | COMMENT |
| --- | --- | --- |
| EXPERIMENTAL DESIGN |  |  |
| Definition of experimental and control groups | Y | In materials and methods |
| Number within each group | Y | Specified in figures |
| SAMPLE |  |  |
| Description | Y | In materials and methods |
| Microdissection or macrodissection | NA |  |
| Processing procedure | Y | In materials and methods |
| If frozen - how and how quickly? | Y | In materials and methods |
| If fixed - with what, how quickly? | Y | In materials and methods |
| Sample storage conditions and duration (especially for FFPE samples) | Y | In materials and methods |
| NUCLEIC ACID EXTRACTION |  |  |
| Procedure and/or instrumentation | Y | In materials and methods |
| Name of kit and details of any modifications | Y | In materials and methods |
| Details of DNase or RNAse treatment | Y | According to kit instructions |
| Contamination assessment (DNA or RNA) | Y | In materials and methods |
| Nucleic acid quantification | N | Not performed |
| Instrument and method | NA |  |
| RNA integrity method/instrument | N | Not performed |
| RIN/RQI or Cq of 3' and 5' transcripts | NA |  |
| Inhibition testing (Cq dilutions, spike or other) | Y | In materials and methods |
| REVERSE TRANSCRIPTION |  |  |
| Complete reaction conditions | Y | In materials and methods |
| Amount of RNA and reaction volume | Y | In materials and methods |
| Priming oligonucleotide (if using GSP) and concentration | NA |  |
| Reverse transcriptase and concentration | Y | According to kit instructions |
| Temperature and time | Y | In supporting information |
| qPCR TARGET INFORMATION |  |  |
| Sequence accession number | N | Not provided |
| Amplicon length | Y | In materials and methods |
| *In silico* specificity screen (BLAST, etc) | NA |  |
| Location of each primer by exon or intron (if applicable) | NA |  |
| What splice variants are targeted? | NA |  |
| qPCR OLIGONUCLEOTIDES |  |  |
| Primer sequences | Y | In materials and methods |
| Probe sequences | Y | In materials and methods |
| Location and identity of any modifications | NA |  |
| qPCR PROTOCOL |  |  |
| Complete reaction conditions | Y | In materials and methods |
| Reaction volume and amount of cDNA/DNA | Y | In materials and methods |
| Primer, (probe), Mg++ and dNTP concentrations | Y | According to kit instructions |
| Polymerase identity and concentration | Y | According to kit instructions |
| Buffer/kit identity and manufacturer | Y | In materials and methods |
| Additives (SYBR Green I, DMSO, etc.) | Y | According to kit instructions |
| Complete thermocycling parameters | Y | In materials and methods |
| Manufacturer of qPCR instrument | Y | In materials and methods |
| qPCR VALIDATION |  |  |
| Specificity (gel, sequence, melt, or digest) | N | Performed but not reported |
| For SYBR Green I, Cq of the NTC | N | Performed but not reported |
| Standard curves with slope and y-intercept | Y | In supporting information |
| PCR efficiency calculated from slope | Y | In materials and methods |
| r2 of standard curve | Y | In materials and methods |
| Linear dynamic range | Y | In materials and methods |
| Cq variation at lower limit | Y | In materials and methods |
| Evidence for limit of detection | Y | In materials and methods |
| DATA ANALYSIS |  |  |
| qPCR analysis program (source, version) | Y | In materials and methods |
| Cq method determination | Y | In materials and methods |
| Outlier identification and disposition | NA |  |
| Results of NTCs | N | Performed but not reported |
| Justification of number and choice of reference genes | NA |  |
| Description of normalisation method | NA |  |
| Number and concordance of biological replicates | Y | Specified in figures |
| Number and stage (RT or qPCR) of technical replicates | Y | In materials and methods |
| Repeatability (intra-assay variation) | N | Not performed |
| Statistical methods for result significance | Y | In materials and methods |
| Software (source, version) | Y | In materials and methods |

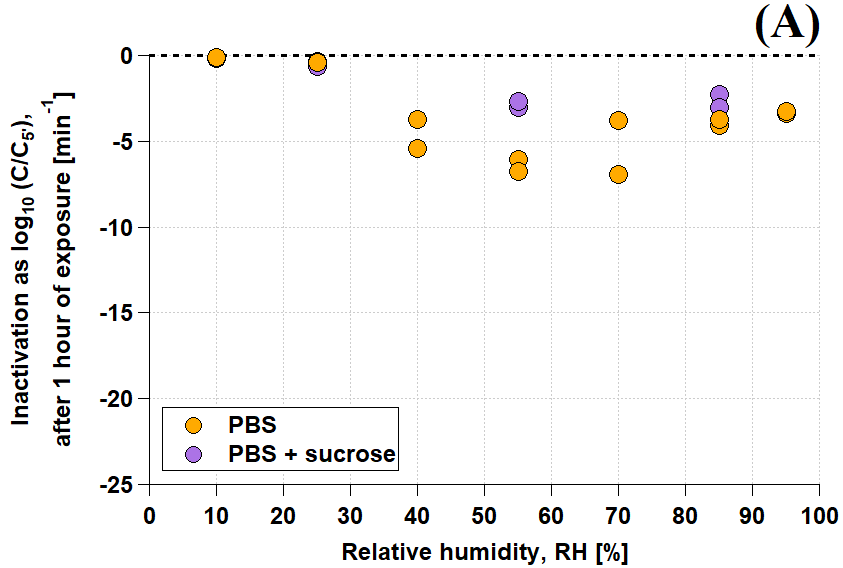

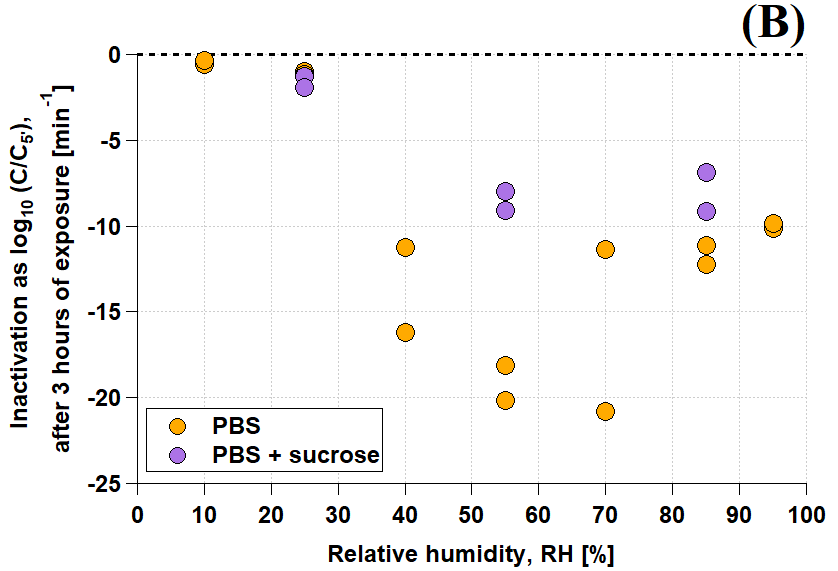

Figure S1. IAV inactivation rate during the aerosol particle experiments performed in the LAPI BREATH (shown as *t*_99_ in Figure 3) after **(A)** 1 hour and **(B)** 3 hours of exposure.

**Potential reasons for the discrepancy between aerosol particle and microliter droplet experiments**

Figure 3 shows that IAV inactivation in 1-μL droplet and bulk experiments by Yang et al. (2012) and Schaub et al. (2023) using saline media is faster by up to an order of magnitude than inactivation in aerosol particles at medium and high RH. This cannot only be explained in terms of size effects. The equilibration between RH in the gas phase and *a_w_* in the liquid phase, i.e., *a_w_* = RH, is virtually instantaneous in submicron aerosol particles, whereas it typically takes less than 30 minutes in 1-μL droplets (Schaub et al., 2023). This could lead to differences of at most a factor of 2 (which has been correctly taken into account in the 1-μL droplet modeling). Furthermore, at RH > 50 %, there should be no efflorescence and potential complications resulting from the formation of complex dendritic morphologies can be avoided. Therefore, the observed large differences are surprising.

As mentioned in the main text of this paper, we can only speculate about the reasons for the discrepancies in *t*_99_. Reasons could be the dependence of *t*_99_ on (i) the initial virus titer, (ii) the pH of PBS, and (iii) exposure to air and strong surface tension forces that damage the viruses in/on the small aerosol particles but not in the large droplets. In the following, we qualitatively discuss some aspects of these potential reasons:

(i)	**Dependence of *t*_99_ on the initial virus titer.** A virus can be affected by the presence of other viruses of the same strain, because they lead to the increase of organic molecules, which are known to have a protective effect (e.g., Kormuth et al., 2018), and possibly because viruses agglomerate. While the initial titers in the bulk and droplet experiments were as high as 10^9^ PFU/mL, we expect that aerosol particles in the LAPI BREATH will typically contain only one virus, if any, based on the work of Zuo et al. (2013) and Pan et al. (2019). To investigate whether this leads to different protection effects, we performed measurements in pure salt solutions with initial titers between less than 10^6^ PFU/mL and more than 10^9^ PFU/mL (see Figure S2). The corresponding *t*_99_ increases by a factor of 4 in this range of titer (as indicated by the slopes of the linear regression lines). We repeated these measurements for 0.14 M NaCl solution (in equilibrium with RH = 99.4 %) and found an even larger enhancement factor (not shown). Therefore, we hypothesize that a factor of 4 within the discrepancy in *t*_99_ might be due to differences in the initial titers between aerosol and large-volume measurements. This is illustrated by the yellow-shaded range in Figure S3.

(ii)	**The dependence of *t*_99_ on the pH of PBS.** The pH value of a pure NaCl solution droplet stays constant during the drying process, namely close to pH 7. In contrast, in a PBS droplet, the partitioning between the ions (Na^+^, Cl**^–^**, H^+^, H_2_PO_4_**^–^**, HPO_4_^2^**^–^**) and their activity coefficients changes continuously during drying, which changes their pH. We measured the pH in PBS solutions at various concentrations and compared with our Pitzer ion interaction model (Luo et al., 2022) and found for 1 ´ PBS (*a_w_* = 0.994) pH = 7.43 (measured) or pH = 7.39 (modeled) and for 20 ´ PBS (*a_w_* = 0.898) pH = 6.58 (measured) or pH = 6.53 (modeled), corresponding to pH-induced reduction in *t*_99_ of about a factor of 3. Therefore, we hypothesize that with continuing drying of the PBS droplets, a growing fraction of the discrepancy in *t*_99_ between aerosol and large-volume measurements might be due to the concomitant pH reduction. This is illustrated by the red-shaded range in Figure S3.

(iii)	**The dependence of *t*_99_ on tension forces.** Finally, we hypothesize that the viruses present in the aerosol particles (typically 100 nm in size), being exposed to air, develop triplet interface lines between the saline liquid, the gas phase and the organic phase of virus, and could therefore experience much stronger forces than in large droplets. This is due to higher surface tension effects on viruses, which are only partially covered by the saline solution, compared to complete immersion in the case of droplets. The surface tension on viruses contained within evaporating droplets may disrupt their envelope and inactivate them (Coleman et al., 2024). The same study concludes that osmotic pressure from dissolved salts still may dominate inactivation of viruses, not because of differences in their magnitude, but rather the exposure time to each force. Further efforts are needed to judge the importance of these effects.

From Figure S3 it is clear that the effects outlined in (i) to (iii) may explain the observed discrepancies to a large degree. However, without additional work, they remain speculative.

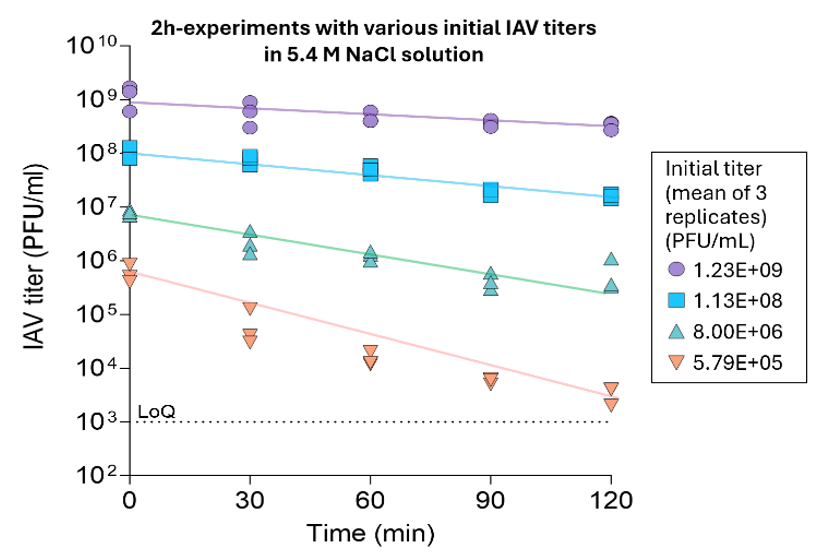

**Figure S2.** **Infective IAV titer evolution in 5.4-M NaCl solution for different initial titers.** Initial virus concentrations are indicated in the box on the right. The molarity of 5.7 M corresponds to a saturated solution, which is in equilibrium with RH = 73 % at room temperature.

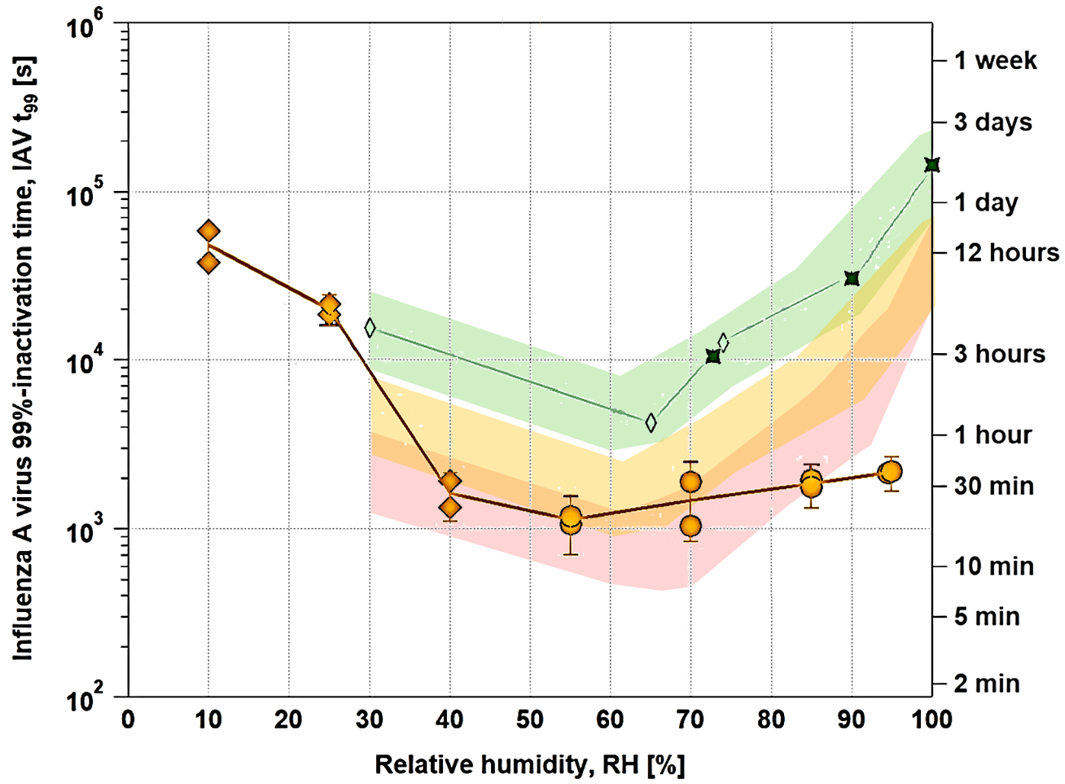

**Figure S3.** **Potential effects of high titer (yellow range) and pH dependence (red range), which might bring bulk and droplet measurements (green range) into agreement with aerosol measurements (orange circles).** Processed RH-dependent IAV infectivity data in PBS from the LAPI BREATH (orange circles) in comparison with our pure NaCl solution measurements in bulk (green stars) and in 1-μL droplets (green diamonds). The high titer in the bulk experiments may explain a factor of 4 of the discrepancy (from green range to yellow range) and the dependence of pH an additional RH-dependent fraction of the discrepancy (from yellow range to red range).
